## Supplementary Figures for "Paradoxical dominant negative activity of an immunodeficiency-associated activating *PIK3R1* variant"

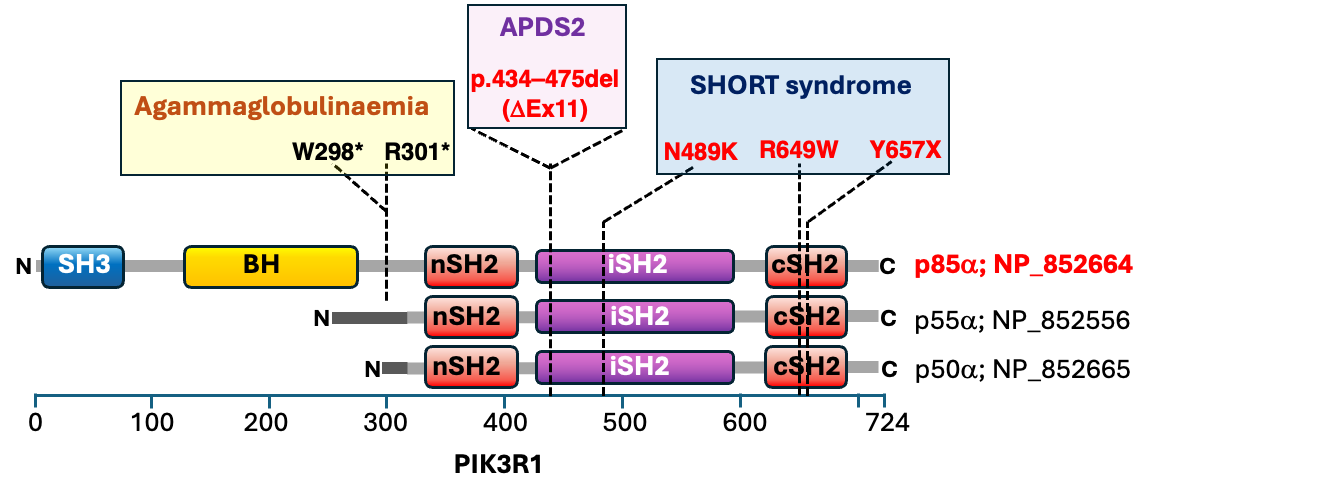

**Figure 1 figure supplement 1. Schematic illustrating the PIK3R1 variants studied.** All three protein products of PIK3R1 are illustrated, namely p85α, p55α and p50α. The site of the heterozygous in-frame deletion caused by skipping of exon 11 that explains most cases of APDS2 is indicated as well as the three heterozygous SHORT syndrome causal variants studied, including the commonest causal variant R649W. All proteins and variants studied are indicated in red. For reference the reported truncating homozygous variants that disrupt only p85α and that are associated with agammaglobulinaemia are also shown. BH = BCR homology, nSH2 = N-terminal SH2, cSH2 = C-terminal SH2, iSH2 = inter-SH2 domain.

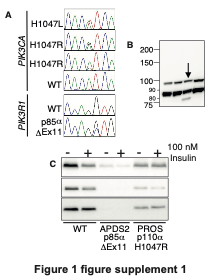

**Figure 1 figure supplement 2. Further characterisation of primary dermal fibroblasts studied.** (A) Details of cDNA sequence for *PIK3CA* and *PIK3R1* from cells derived from healthy controls (WT), patients with APDS2 (p85α ΔEx11) or *PIK3CA*-related overgrowth syndrome (PROS), confirming expected expression of mutant alleles. (B) Higher magnification detail of immunoblot from wild-type and APDS2 fibroblasts (arrowed lane) showing truncated p85α Δex11 in APDS2 cells only. (C) Close up view of all 3 immunoblot replicates for p110δ blots quantified in Figure 1 (APDS2 and adjacent genotypes only), showing severely reduced p110δ expression in the APDS2 cell line

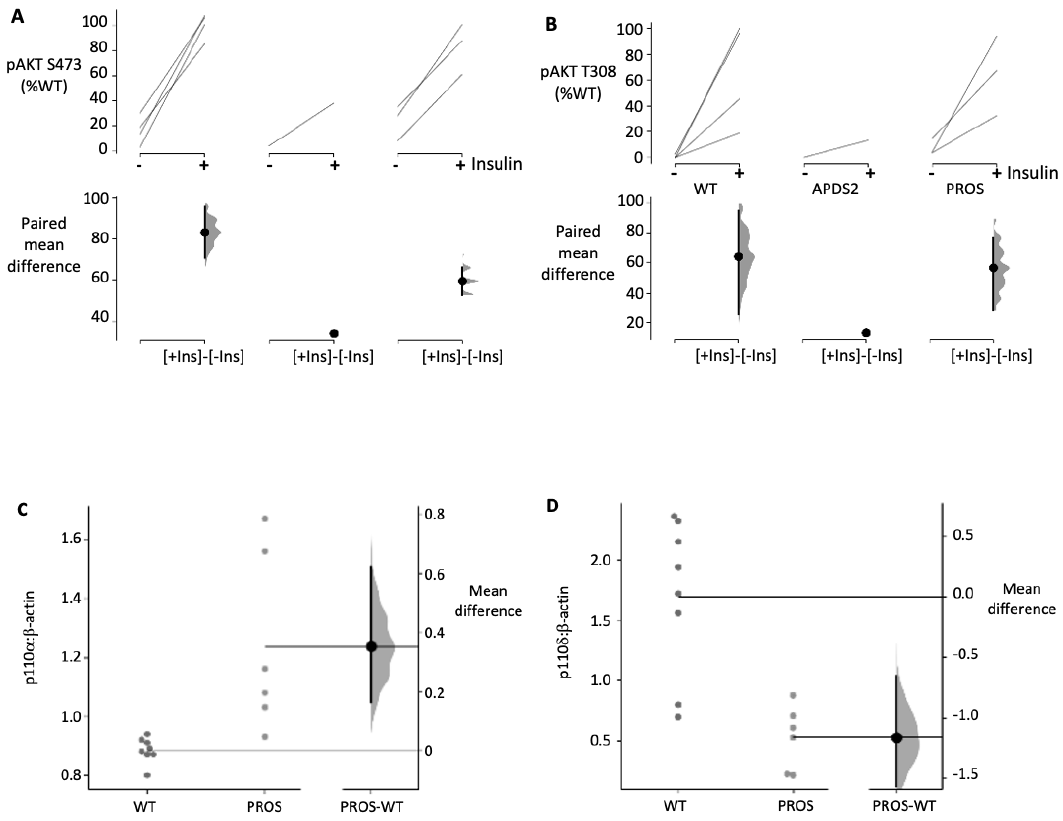

**Figure 1 figure supplement 3. Full statistical analysis of data presented in main Figure 1** Analysis of insulin-induced increase in (A) AKT S473/4 and (B) T308/9 phosphorylation. The paired mean difference for 3-4 comparisons are shown in Cumming estimation plots. The raw data, as presented in Figure 1, are re-plotted on the upper axes with paired observations connected by a line. On the lower axes, paired mean differences are plotted as a bootstrap sampling distribution. Mean differences are depicted as dots; 95% confidence intervals are indicated by the ends of the vertical error bars. (C-D) Analysis of differences in (C) p110α and (D) p110δ protein expression between healthy control cells and cells from PROS patients harbouring activating PIK3CA mutations. Mean differences are shown in Gardner-Altman estimation plots, with expression data plotted on the left axes and mean difference on floating axes on the right, again as a bootstrap sampling distribution with mean difference depicted as a dot and 95% confidence intervals by the ends of the vertical bar.

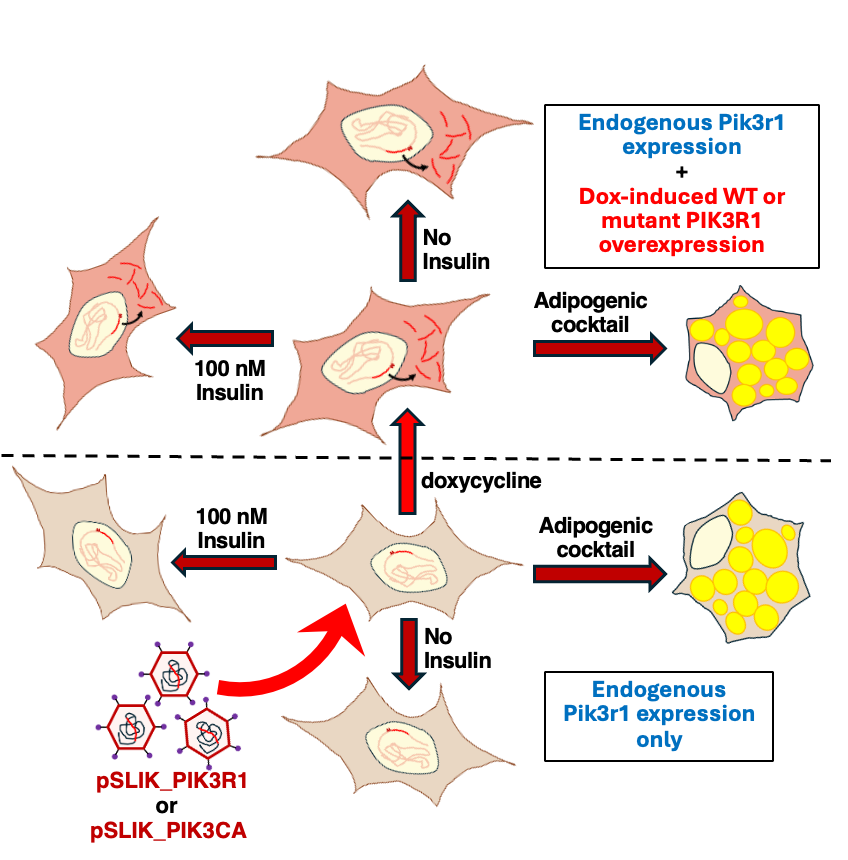

### Figure 2 figure supplement 1. Schematic illustrating experimental design for 3T3-L1 studies. Wild type 3T3-L1 murine preadipocytes with intact endogenous Pik3r1 expression were infected with pSLIK lentivirus with a payload of wild-type or mutant human PIK3R1 under control of a doxycycline-responsive promoter. After selection stable cells with or without doxycycline exposure to induce transgenic PIK3R1 expression were either differentiated to adipocytes, or stimulated with 100nM insulin as indicated.

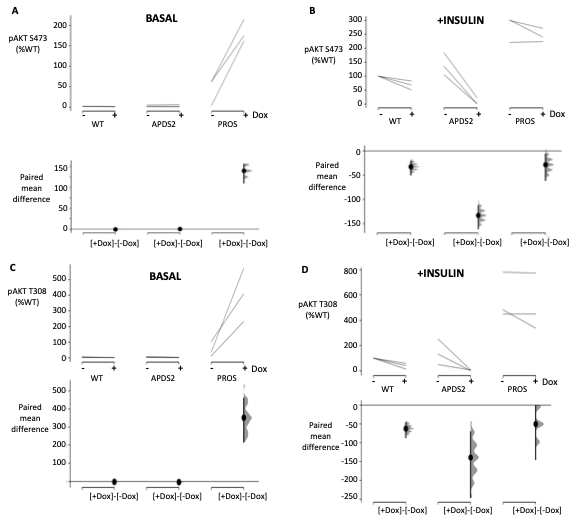

### Figure 2 figure supplement 2. Full statistical analysis of data presented in main Figure 2. Analysis of the effects of doxycycline-induced expression of wild-type (WT) or ΔEx11 (APDS2) p85α, or of p110α H1047R (PROS) on Akt S473/4 (A,B) and T308/9 (C,D) phosphorylation. Comparisons are made in both the basal, non-insulin stimulated state (A,C) and after stimulation with 10 nmol/L insulin (B,D). Paired mean differences for 3 comparisons are shown in Cumming estimation plots. Raw data, as presented in Figure 2, are re-plotted on the upper axes with paired observations connected by a line. On the lower axes, paired mean differences are plotted as a bootstrap sampling distribution. Mean differences are depicted as dots; 95% confidence intervals are indicated by the ends of the vertical error bars.

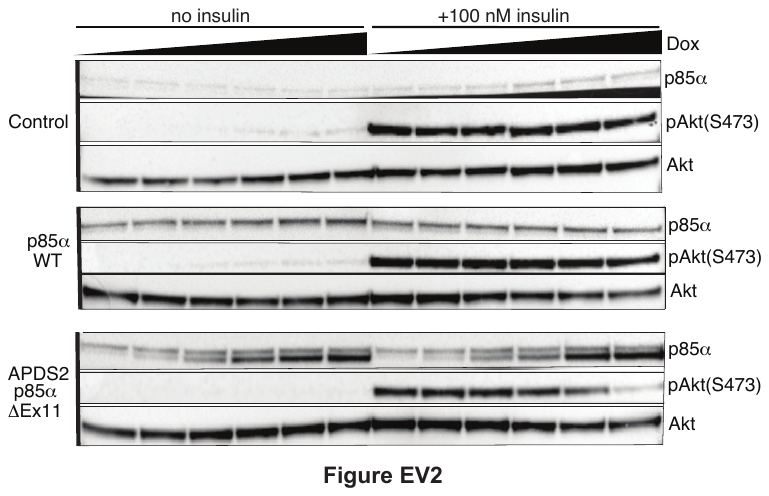

**Figure 2 figure supplement 3. The effect of graded expression of wild-type or disease-associated p85α on 3T3-L1 preadipocytes.** Immunoblots of p85α, phosphoAkt (S473) and total Akt are shown for control 3T3-L1 cells and 3T3-L1 cells conditionally expressing wild-type (WT), or APDS2-associated mutant p85α under the control of doxycycline (Dox), with and without 10 minutes of exposure to insulin as indicated. The filled black triangles indicate increasing concentrations of doxycycline (from left to right: 0, 0.02, 0.03, 0.045, 0.065, or 0.1 μg/mL). Exposure was for 72 hours in all cases. The truncated p85α variant can be seen below the WT p85α for the APDS2 ΔEx11 mutant.

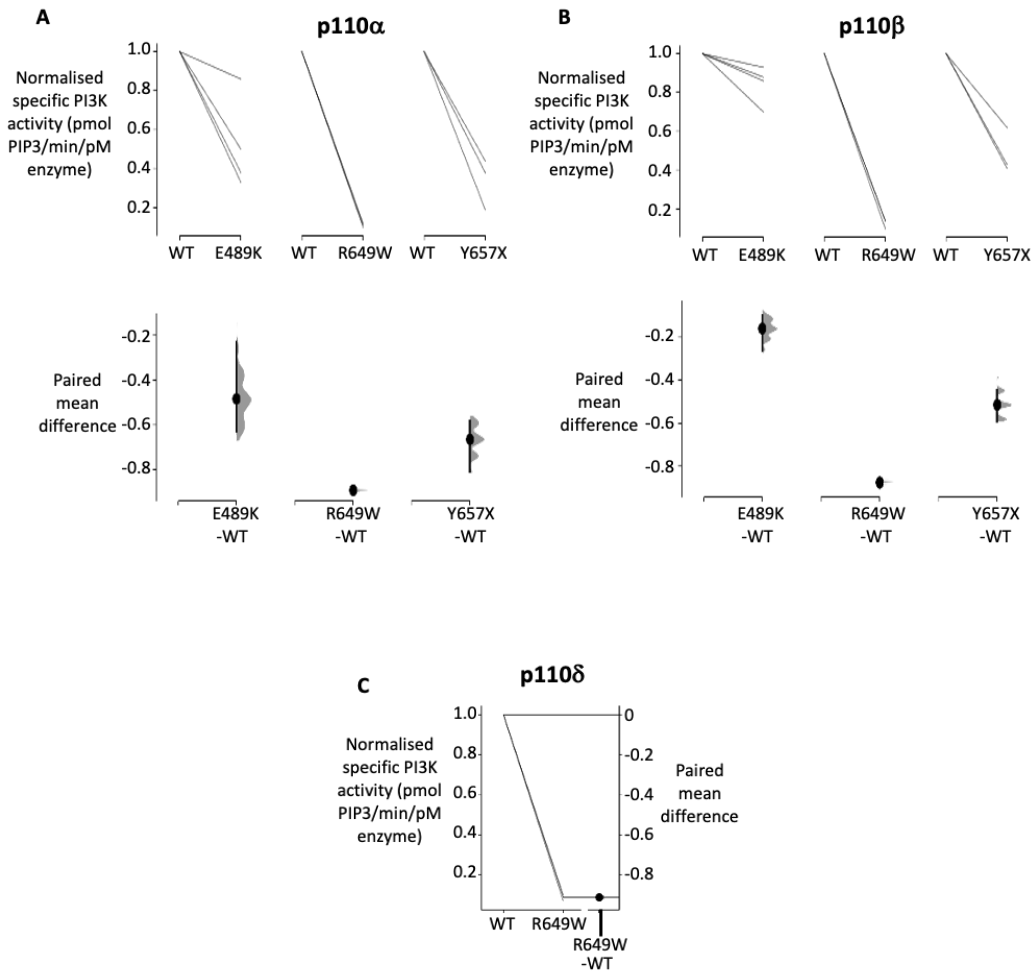

**Figure 3 figure supplement 1. Full statistical analysis of data presented in main Figure 3.** Analysis of fluorescence polarisation assay of phosphoinositide 3-kinase (PI3K) activity of *in vitro* synthesised wild-type (WT) or mutant (E489K, R649W or Y657X) p85α. Results for p110α- and p110β-containing PI3K are shown in (A) and (B) respectively. All data were acquired in the presence of phosphotyrosine peptide. Paired mean differences for 3 comparisons are shown in Cumming estimation plots. Raw data, as presented in Figure 3, are re-plotted on the upper axes with paired observations connected by a line. On the lower axes, paired mean differences are plotted as a bootstrap sampling distribution. Mean differences are depicted as dots; 95% confidence intervals are indicated by the ends of the vertical error bars. Results for the R649W p85α mutation only are shown with p110δ in (C). In this case raw data are re-plotted on the left hand axes with paired observations connected by 3 nearly superimposed lines. On the right hand axes, paired mean differences are plotted as a bootstrap sampling distribution.

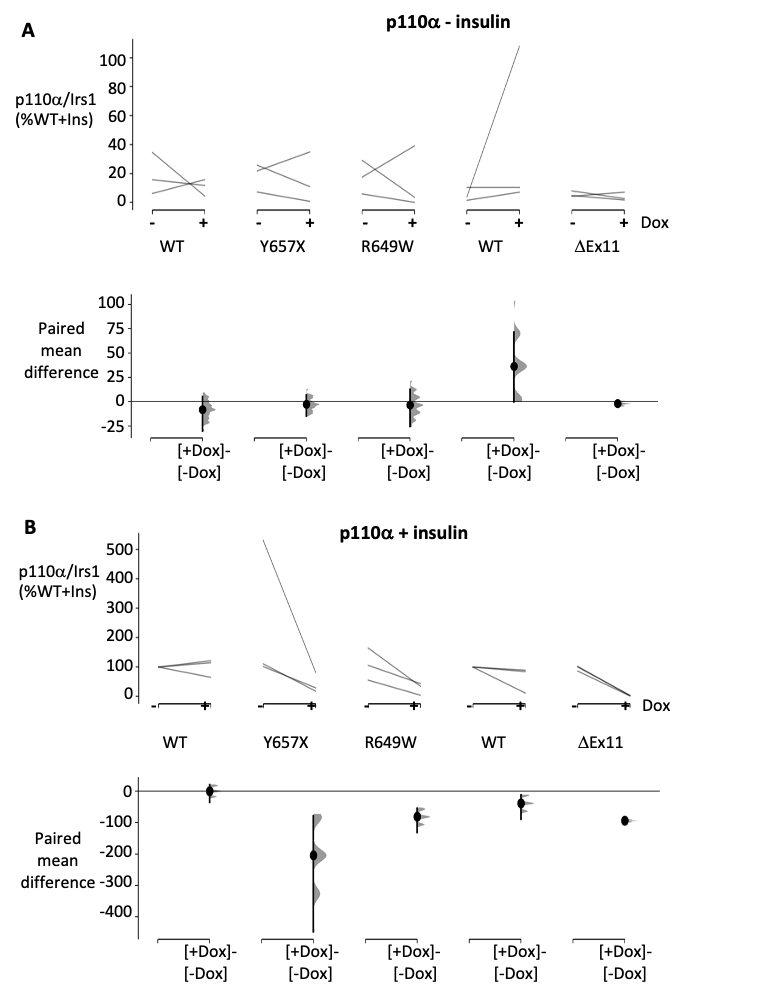

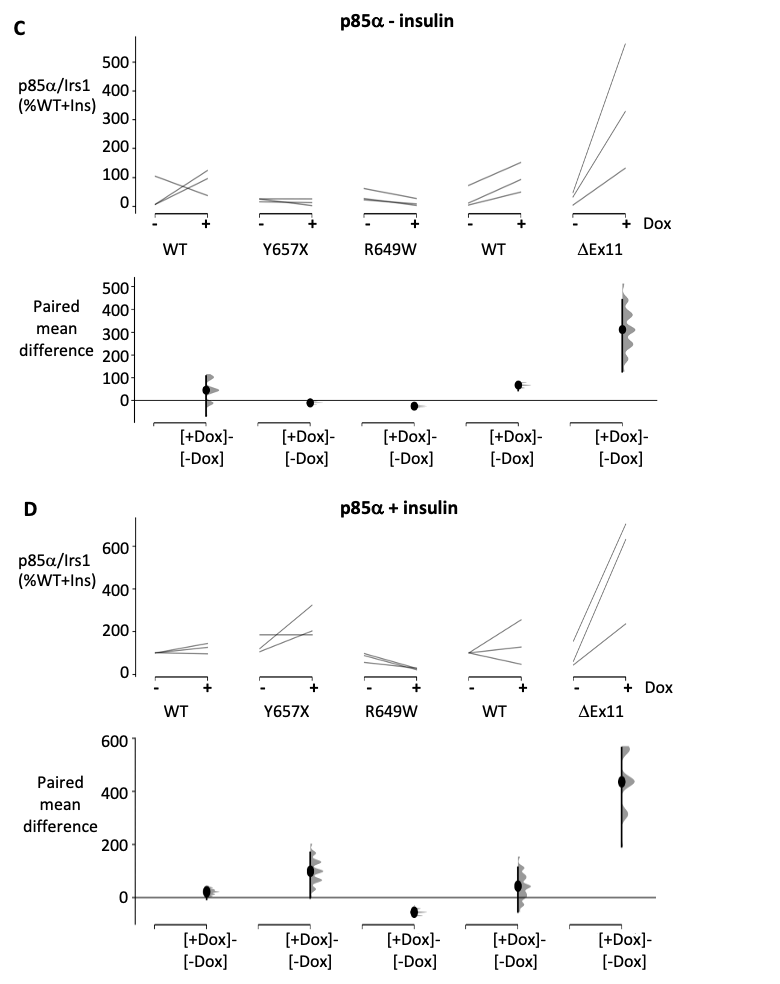

### Figure 5 figure supplement 1. Full statistical analysis of data presented in main Figure 5. Analysis of the effects of doxycycline (dox)-induced expression of wild-type (WT), Y657X, R649W or ΔEx11 p85α on association of p110α (A,B) and p85α (C,D) with Irs1. Results of co-immunoprecipitation with (B,D) and without exposure to 10 nmol/L insulin Comparisons are made in both the basal, non-insulin stimulated state (A,C) and after stimulation with 10 nmol/L insulin (B,D) are shown. Paired mean differences for 3 comparisons are shown in Cumming estimation plots. Raw data, as presented in Figure 5, are re-plotted on the upper axes with paired observations connected by a line. On the lower axes, paired mean differences are plotted as a bootstrap sampling distribution. Mean differences are depicted as dots; 95% confidence intervals are indicated by the ends of the vertical error bars.

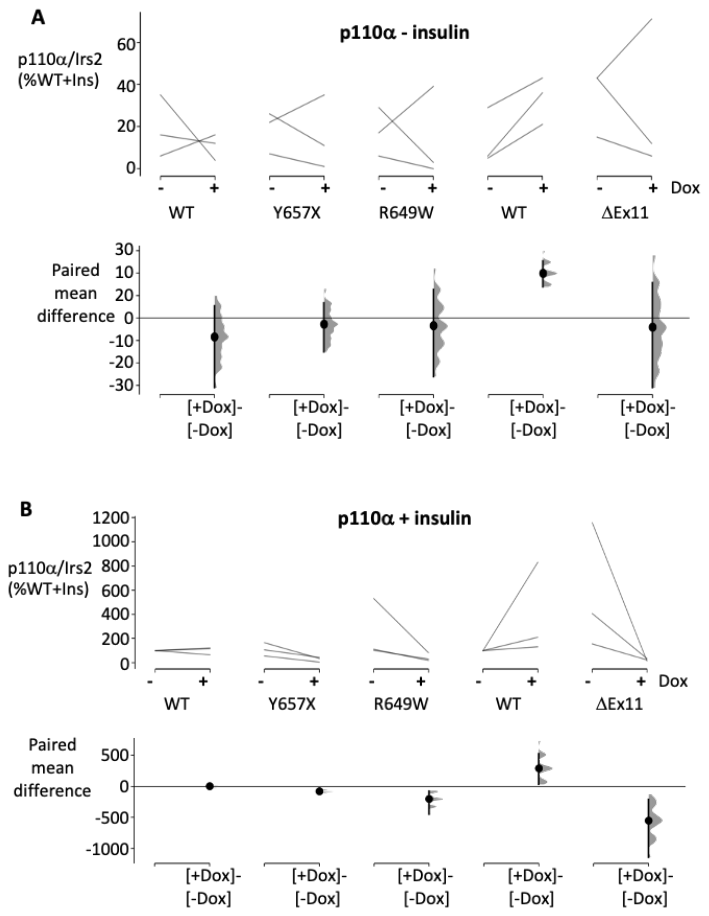

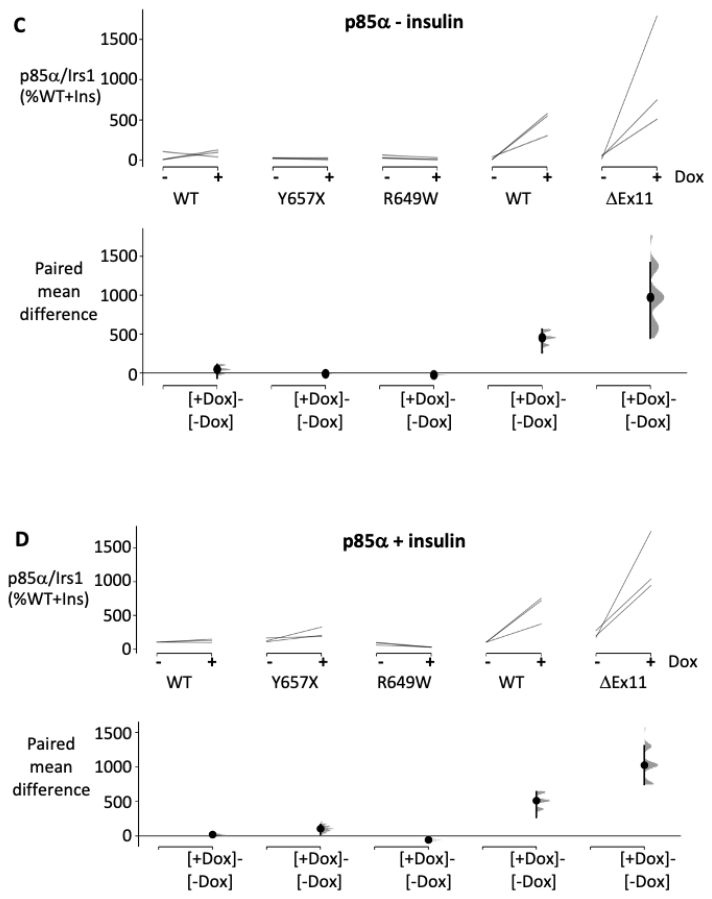

### Figure 6 figure supplement 1. Full statistical analysis of data presented in Figure 6. Analysis of the effects of doxycycline (dox)-induced expression of wild-type (WT), Y657X, R649W or ΔEx11 p85α on association of p110α (A,B) and p85α (C,D) with Irs2. Results of co-immunoprecipitation with (B,D) and without exposure to 10 nmol/L insulin Comparisons are made in both the basal, non-insulin stimulated state (A,C) and after stimulation with 10 nmol/L insulin (B,D). Paired mean differences for 3 comparisons are shown in Cumming estimation plots. Raw data, as presented in Figure 6, are re-plotted on the upper axes with paired observations connected by a line. On the lower axes, paired mean differences are plotted as a bootstrap sampling distribution. Mean differences are depicted as dots; 95% confidence intervals are indicated by the ends of the vertical error bars.
